# Spared nerve injury induces long-term and brain region-specific changes in oligodendrocyte density in mice

**DOI:** 10.1101/2025.10.01.679851

**Authors:** Léa J. Becker, Rory O’Shea, Gustavo Borges, Jordan G. McCall

**Affiliations:** Department of Anesthesiology, Center for Clinical Pharmacology, Washington University Pain Center, Washington University in St. Louis, St. Louis, MO, USA

## Abstract

**Background:** Emerging evidence suggests a role for non-neuronal cells in the pathophysiology of chronic pain. Chronic pain causes profound alterations of the transcriptomic program of oligodendroglia, but the effect of pain on the oligodendroglial cells themselves remains unknown.

**Methods:** Male and female C57BL6/J mice underwent spared nerve injury (SNI). Mechanical hypersensitivity was assessed five weeks later using the von Frey test. Six weeks post-surgery, mice were perfused, brains dissected, and immunostained for oligodendrocyte precursors cells (OPC) and mature oligodendrocytes (OL) in four brain regions involved in pain chronification: anterior cingulate cortex (ACC), central amygdala (CeA), basolateral amygdala (BLA), and periaqueductal gray (PAG).

**Results:** We found OL were reduced in the ACC and CeA of both sexes six weeks after SNI. Conversely, BLA OL were increased in both sexes following SNI. There was a sex-dependent effect of SNI on PAG-OL, where OL were only reduced in females. SNI did not affect OPC in any of the studied brain regions, but female PAG and BLA appeared to have fewer OPC than males independent of SNI.

**Conclusion:** Long-term nerve injury differentially affects OL in a brain region- and sex-dependent manner. This effect is observable six weeks after injury, suggesting a long-lasting impact of chronic pain on oligodendroglial cells. OPC, on the other hand, are remarkably stable. This finding aligns with previous literature showing OPC maintain homeostasis, even in pathological conditions.

**Significance:** Oligodendrocytes ensheathe axons to increase conduction speed, stabilize neuronal connections, and fine-tune neuron-to-neuron communication. Furthermore, pharmacological stimulation of myelination improves pain-induced cognitive deficits in mice, suggesting therapeutic potential in targeting oligodendrocytes in pain. A better understanding of how pain impacts oligodendroglia is thus crucial to better understand the pathophysiology of pain and identify new therapeutic targets.

## 1. Introduction

Chronic pain is a debilitating disease that affects approximately 20% of the global population (Rikard et al., 2023; Zimmer et al., 2022). Despite its severe social and economic burden, current treatments for chronic pain often remain inadequate, urging the need to identify new therapeutic targets. Although neurons have been the main focus of chronic pain research, a growing body of evidence now underscores the importance of non-neuronal cells (Lee et al., 2023; Marty-Lombardi et al., 2024; Tansley et al., 2022). For example, astrocytes and microglia are implicated in chronic pain-induced central nervous system (CNS) inflammation (Li et al., 2022; Ni et al., 2016; Vergne-Salle and Bertin, 2021), and damage to myelinating Schwann cells in the periphery is a well-established feature of peripheral neuropathy (De Logu et al., 2017; Hartlehnert et al., 2017). Surprisingly, the impact of pain on oligodendrocytes, the source of myelination in the CNS, is less documented (Kim and Angulo, 2025; Malta et al., 2019), despite the high comorbidity between CNS-demyelinating diseases and pain (Asseyer et al., 2021; Diaz et al., 2025; Ichimiya et al., 2023; Koc et al., 2025; Potter et al., 2016). Thus, gaining insights into how cells of the oligodendrocyte lineage are affected by chronic pain may uncover new pain relief mechanisms.

The oligodendrocyte lineage comprises several cellular subtypes, whose classification is still being refined, but can generally be divided into two main cell types: oligodendrocytes (OL) and oligodendrocyte precursor cells (OPC) (Almeida and Lyons, 2017; Marques et al., 2019, 2016; Spitzer et al., 2019). OL are primarily responsible for ensheathing axons in the CNS, leading to faster signal conduction, increased metabolic support for neurons, and stabilized and fine-tuned neuronal connections (Almeida and Lyons, 2017; Mount and Monje, 2017; Saab and Nave, 2017). OPC are the most proliferative cells in the brain, even in adulthood (Richardson et al., 2011). A small subset of OPC will differentiate into mature OL, while others will continue to proliferate and regulate important physiological processes, including ion homeostasis, immune responses, and phagocytosis (Buchanan et al., 2023; Dimou and Gallo, 2015; Kahng et al., 2025; Maldonado et al., 2013). OPC also respond to neuronal activity through direct synaptic inputs (Bergles et al., 2010; Nagy et al., 2017). Both OPC and OL have been implicated in the pathophysiology of chronic pain, mostly at the transcriptomic level (Becker et al., 2023; Dai et al., 2022). A recent study showed that stimulation of myelination with the anticholinergic antihistamine, clemastine, alleviates chronic pain-induced cognitive deficits in mice (Zhu et al., 2024). However, the precise impact of chronic pain on oligodendrocyte lineage cells remains largely unexplored.

Here, using the mouse spared nerve injury (SNI) model, we examined how OL and OPC cell populations are impacted in brain regions known for their involvement in pain control and chronification: the anterior cingulate cortex (ACC) (Chen et al., 2014; Sellmeijer et al., 2018), central amygdala (CeA) (Kiritoshi et al., 2024; Mazzitelli et al., 2024), basolateral amygdala (BLA) (Apte et al., 2025; Li et al., 2013), and the periaqueductal gray (PAG) (Heinricher et al., 2004; Lubejko et al., 2024; Samineni et al., 2017; Sirucek et al., 2025). We observed that six weeks post-surgery, the number OPC remained unchanged while OLs density was altered in a region-specific fashion.

## 2. Methods

### 2.1 Animals

All experiments were conducted using male and female adult C57BL/6J mice (JAX:000664), 10-13 weeks old at the beginning of experimental procedures. Mice were originally sourced from The Jackson Laboratory (Bar Harbor, ME, USA) and bred in-house in a barrier facility in another building before being transferred to a holding facility adjacent to the behavioural space between 4-6 weeks of age. Mice were left undisturbed except for necessary husbandry to habituate to the new facility until 10 weeks of age. All mice were group-housed, given *ad libitum* access to standard laboratory chow (PicoLab Rodent Diet 20, LabDiet, St. Louis, MO, USA) and water, and maintained on a 12:12-hour light/dark cycle (lights on at 7:00 AM without shifting for daylight savings time). All experiments and procedures were approved by the Institutional Animal Care and Use Committee of Washington University School of Medicine in accordance with National Institutes of Health and ARRIVE guidelines.

### 2.2 Spared nerve injury (SNI)

SNI was used to induce neuropathic injury and performed as previously described (Cichon et al., 2018; Norris et al., 2024a). Mice were assigned to experimental groups prior to surgery to ensure that groups did not initially differ in mechanical withdrawal threshold. Mice were anesthetized with 3% isoflurane. The right hind limb was shaved and disinfected with 75% ethanol and betadine. A 10-15 mm incision was made in the skin proximal to the knee to expose the biceps femoris muscle. Separation of the muscle allowed visualization of the sciatic nerve trifurcation. The common peroneal and tibial branches were ligated with 6-0 silk suture (Ethicon Inc., Raritan, NJ, USA) and 1 mm of nerve was excised distal to the ligature, leaving the sural branch intact. Following wound closure mice were allowed to recover on a table warmed to 43°C for 10-20 minutes prior to being returned to their home cage. Sham surgeries were identical to the SNI procedure without the ligation, excision, and severing of the peroneal and tibial branches of the sciatic nerve.

### 2.3 von Frey filament test

Mechanical thresholds of the hind paws were evaluated using von Frey hairs (Bioseb, Pinellas Park, FL,USA) (Becker et al., 2023; Benbouzid et al., 2008; Yalcin et al., 2014). Mice were placed in clear Plexiglas® boxes (12 cm diameter) on an elevated mesh grid and left to habituate for 15 minutes. Filaments were applied to the plantar surface of each hind paw in a series of ascending forces (0.16–6 g). Each filament was tested five times on the lateral portion of each paw until it just bent. The threshold was defined as three or more withdrawals observed out of five trials. All animals were tested twice: once before SNI surgery to determine the basal threshold and once five weeks after surgery to ensure the development of mechanical hypersensitivity (**Fig. 1a-b**). Due to guarding of the injured paw in SNI mice, experimenters were unable to be fully blinded during the final von Frey test.

**Figure 1.**
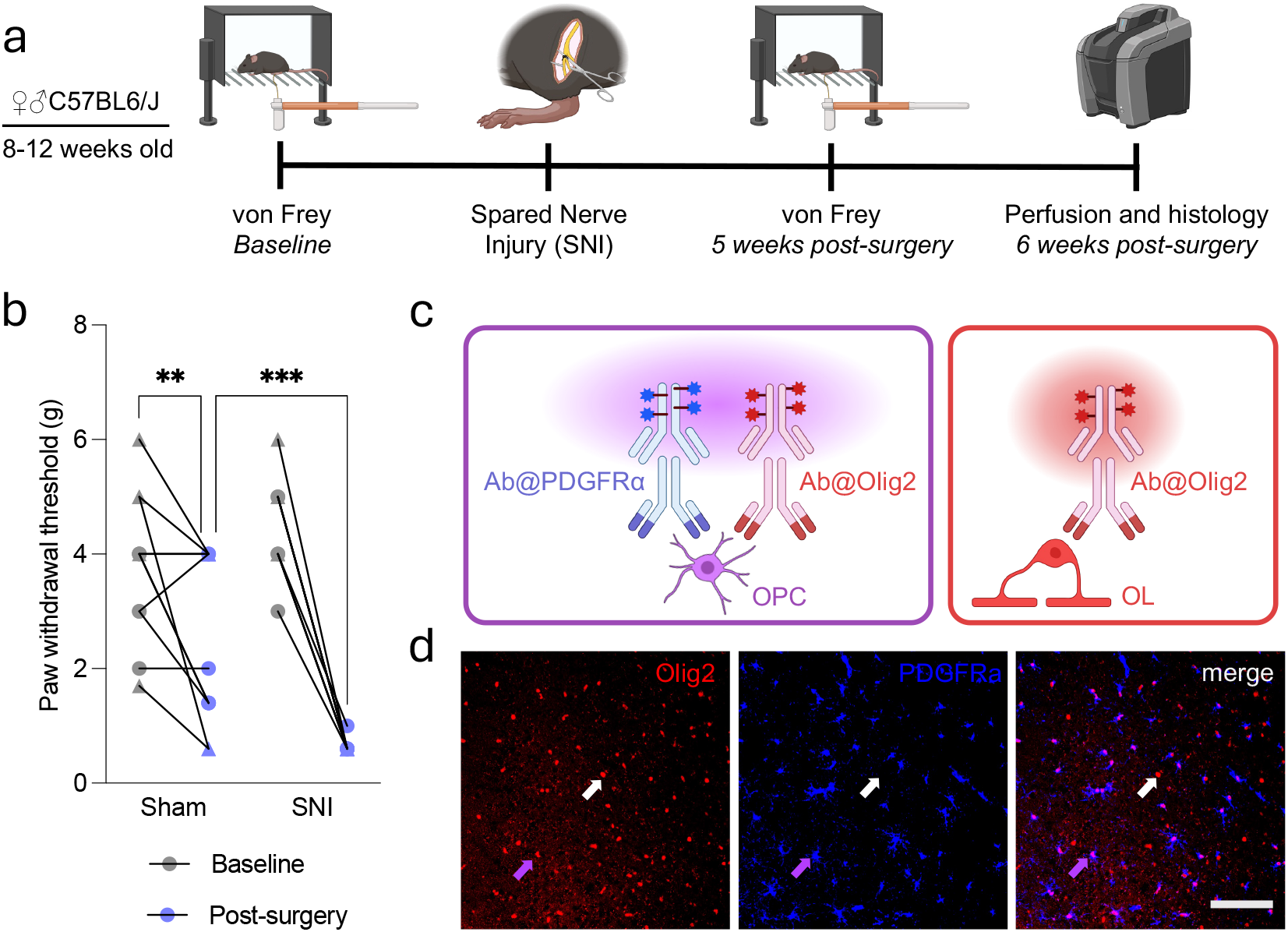
Approach used to assess oligodendrocyte lineage cells following neuropathic injury. (**a**) Experimental design and calendar. (**b**) Five weeks after spared nerve injury (SNI) mice show decreased paw withdrawal thresholds in the hind paw ipsilateral to the lesion (n_ctrl_=11/n_SNI_=10; circle points = males, triangle = females. (**c**) Immunofluorescence-based approach used to quantify oligodendrocyte precursors (OPC) and mature oligodendrocytes (OL). (**d**) Representative image of stained OPC (purple arrow) and OL (white arrow). Data are represented as mean ± SEM. Scale bar = 100µm.

### 2.4 Tissue preparation and histochemistry

Six weeks after SNI surgery, animals were deeply anesthetized with a cocktail of ketamine/xylazine/acepromazine (69.57 mg/ml; 4.35 mg/ml; 0.87 mg/ml; 182 mg/kg, i.p.) and transcardially perfused with 0.1 M phosphate buffer (PB) followed by ice-cold 4% paraformaldehyde in 0.1 M PB. Brains were dissected, postfixed overnight and kept at 4°C cryoprotected with solution of 30% sucrose in 0.1 M PB at 4°C for at least 24 hours before cutting. 40 µm coronal sections were obtained using a microtome and serially collected in PBS.

Slices containing the ACC, BLA, CeA and PAG were washed in PBS (3×10 min) and pre-incubated in PBS containing Triton X-100 (0.3%) and normal goat serum (5%) for 1 hour. Sections were then incubated overnight at 4°C in PBS containing Triton X-100 (0.3%), rabbit anti-PDGFRa (1:1000, Invitrogen, MA5-41209) and chicken anti-Olig2 (1:500, AvesLab, OLIG2-0020). Sections were then washed in PBS (3 × 10 minutes), incubated with Alexa Fluor 594 goat anti-rabbit secondary antibody (1:500, Invitrogen, A11012) and Alexa Fluor 488 goat anti-chicken (1:500, Invitrogen, A11039) in PBS containing Triton X-100 (0.3%) for 2 hours and washed in PBS (3 × 10 minutes). Finally, sections were serially mounted with Vectashield Hardset with DAPI (Vector Laboratories Inc., H-1500).

### 2.5 PDGFRa and Olig2 quantification

All images were acquired on a BZ-X810 Keyence microscope (Keyence, Itasca, IL, USA) at 10x from 3-4 sections of each structure. For the ACC, four fields of view were stitched together to get a full image of the structure. Positive cells were counted using Fiji. A region-of-interest was defined on the composite image for each brain region, then reported in both red and green channels. A binary image was created using an intensity threshold, and cells were counted separately in the green (Olig2) and red (PDGFRa) channels. The outlines of the counted cells were reported on the initial composite image and the accuracy of the detection was visually verified by a blinded experimenter. OPC were defined as cells stained with both green and red while OL were stained in green only (**Fig. 1c-d**, colors were changed to red (Olig2) and blue (PDGFRa) for accessibility purpose).

### 2.6 Statistics

Statistical analyses were conducted using Prism 10.0 (GraphPad) or R. Von Frey data are expressed as mean ± SEM, and immunohistochemistry as median and quartiles. For immunohistochemistry data, differences between groups were determined using a two-tailed independent Mann-Whitney test, with significance set at p < 0.05. Because measures in the Von Frey test are non-continuous by nature, differences between groups were determined using the nparLD R package with F1-LD-F1 design (Noguchi et al., 2012). Raw data and all statistical tests can be found in **Table S1**.

## 3. Results

Five weeks after SNI, male and female mice had significantly reduced ipsilateral paw withdrawal thresholds (**Fig. 1a-b**, F_(2.744, ∞)_ = 5.875, p=0.0008, post-hoc SNI-ipsilateral_(baseline vs post-surgery)_ <0.0001). Decreased mechanical thresholds were also observed in sham mice, although of lower magnitude (F_(2.744, ∞)_ = 5.875, p=0.0008, post-hoc Sham-ipsilateral_(baseline x post-surgery)_ = 0.0107). This effect was largely driven by females and was possibly due to abnormally high threshold during baseline measurement. No significant difference in paw withdrawal threshold was observed in the non-injured paw of sham mice (F_(2.744, ∞)_ = 5.875, p=0.0008, post-hoc Sham-contralateral_(baseline x post-surgery)_ = 0.0677), but a decrease was detected in the contralateral paw of the SNI group (F_(2.744, ∞)_ = 5.875, p=0.0008, post-hoc SNI-contralateral_(baseline vs post-surgery)_ p = 0.0045), similar to what we observed previously at longer timepoints post-injury (Norris et al., 2024a) (Fig. S1).

Six weeks following SNI, mice presented with several changes in OL density across the assayed brain regions (**Fig. 2a-d**). OL were decreased in both the ACC (**Fig. 2e**, U=12, p=0.0418) and the CeA (**Fig. 2f**, U=16, p=0.0203). Interestingly, however, the BLA showed the opposite response, with OL counts increased following SNI (**Fig. 2g**, U=16, p=0.0097). SNI did not appear to alter the number of OL in the PAG (**Fig. 2h**). However, the effect of SNI on the PAG may be sex-dependent, as a separate analysis, taking sex as a covariate, showed that females had decreased OL density (F_(1,16)interaction_=4.936, p=0.0411, post-hoc females_(sham vs SNI)_ p=0.0431). No effect of SNI was observed on OPC in any of the selected structures (**Fig. 2i-l**). Notably, however, females from both SNI and sham groups had fewer OPC in the BLA and the PAG than males, suggesting these brain regions have basal sex differences in total OPC numbers (BLA_(males vs females)_: F(1, 16)=6.628, p=0.0204, PAG_(males vs females)_: F(1, 16)= 12.73, p=0.0026).

**Figure 2.**
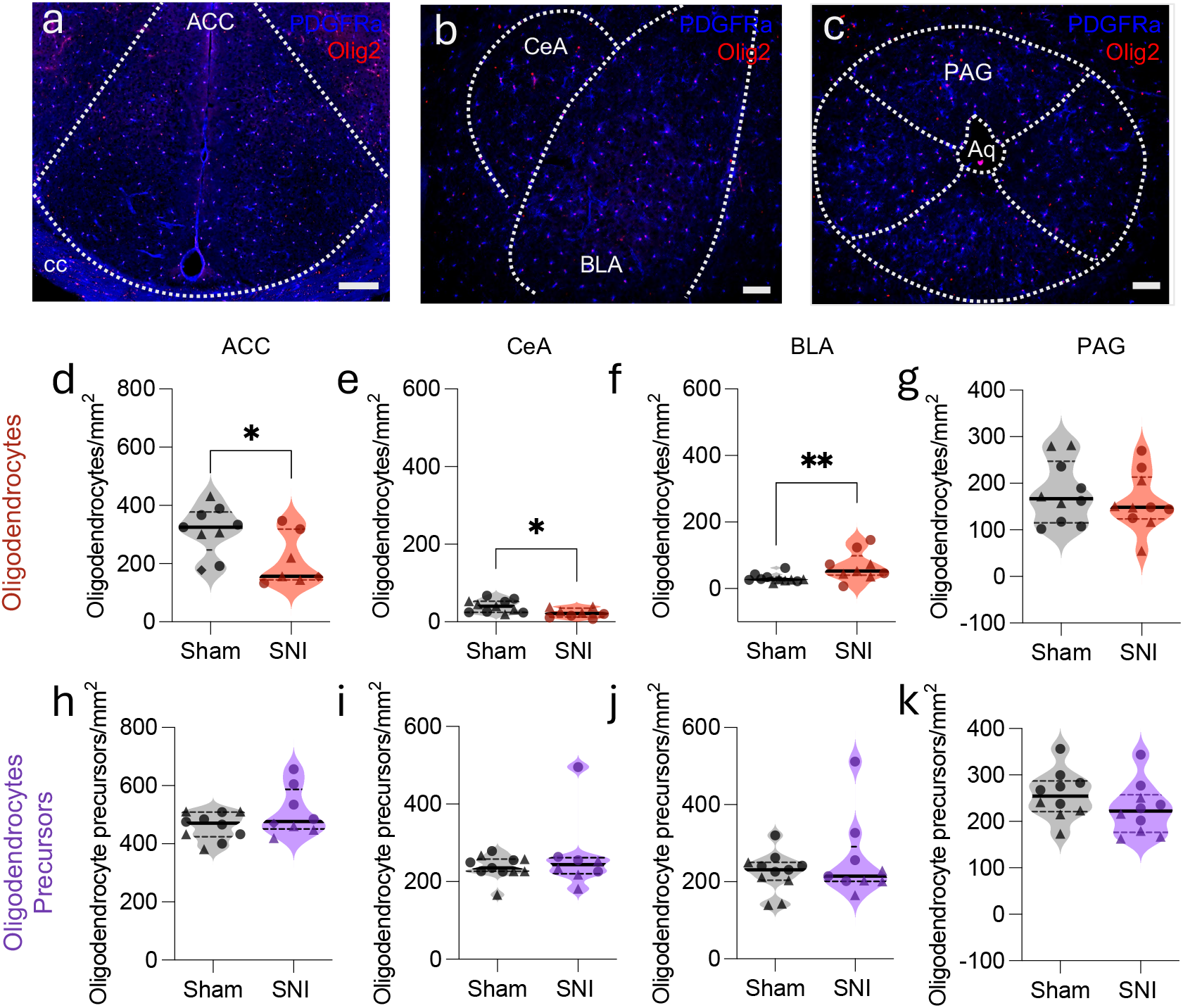
SNI differentially alters OL density in a brain region-dependent manner. Representative images of PDGFRα (blue) and Olig2 (red) staining in the (**a**) ACC, (**b**) CeA and BLA and (**c**) PAG. 6 weeks after SNI surgery the number of OLs is decrease in the (**d**) ACC and the (**e**) CeA, increased in the (**f**) BLA and unaltered in the (**g**) PAG. No effect of SNI was observed in OPC in the ACC, CeA, BLA or PAG (**h-k**). Data are represented as median (solid black line) with 1^st^ and 3^rd^ quartiles (dashed black lines). Circle points = males, triangle = females. Y-axis are scaled by structure. Scale bar = 200µm (ACC) and 100µm (CeA, BLA, PAG). *p<0.05, **p<0.01 n_ctrl_ = 9-11/n_SNI_ = 7-10

We next assessed whether the observed changes in OL count correlate with the severity of SNI-induced mechanical hypersensitivity. In the CeA, a direct correlation between number of OL and hind paw withdrawal threshold was detected (**Fig. 3a**, r^2^=0.2528, p=0.0282). A similar trend was observed in the ACC, although not reaching significance (**Fig. 3b**, r^2^=0.1958, p=0.0861). No correlation was found between mechanical hypersensitivity and the number of OL in the BLA or the PAG (**Fig. S2** r^2^=0.1027, p=0.1683).

**Figure 3.**
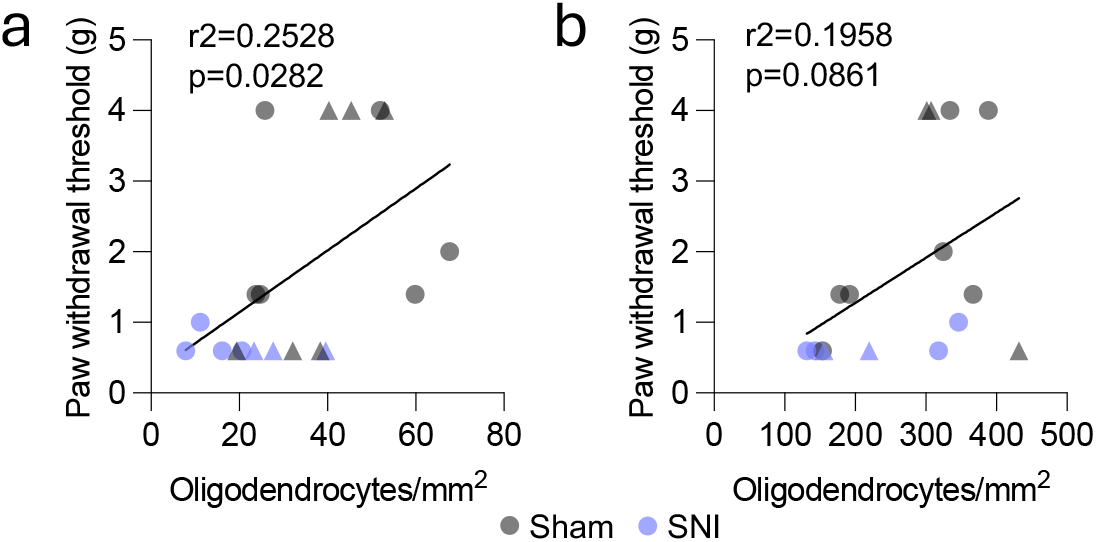
Correlations between OL density and mechanical hypersensitivity. In the CeA (**a**) the number of oligodendrocytes is positively correlated with paw withdrawal threshold in the von Frey test. A similar trend is seen in the ACC (**b**) n_ctrl_ = 10-11/n_SNI_ = 8. Circle points = males, triangle = females.

## 4. Discussion

We show that neuropathic injury leads to long-term changes in OL. This effect varies between brain regions, with decreased OL counts in the ACC and the CeA, but increased OL counts in the BLA. Similar bidirectional alterations have been reported in major depression (Becker et al., 2023; Breton et al., 2021; Lutz et al., 2021, 2017; Zhao et al., 2024). Considering the comorbidity between pain and depression, especially at later time points following neuropathic injury (de la Rosa et al., 2024; Kremer et al., 2021; Norris et al., 2024a; Yalcin et al., 2011), disruption of OL density may represent a convergent target to alleviate pain and its associated negative affect. Furthermore, we also found that chronic injury appears to have a sex-dependent effect on PAG OL, with decreased OL density only present in SNI females. Our study, however, was not initially designed to test for sex differences, and therefore this observation should be tested further in future studies. This is particularly important due to the female preponderance often reported in demyelinating diseases (Diem et al., 2022), possibly linked to intrinsic sexual dimorphism in OL and OPC physiology (Alvarez-Sanchez and Dunn, 2023; Yasuda et al., 2020). Importantly, we only focused on four structures known for their involvement in pain control and chronification. Expanding this investigation to other pain-related CNS regions will be crucial to have a more complete understanding of how OL respond to chronic pain.

In contrast to OL, OPC remained remarkably stable following peripheral neuropathic injury. This may stem from the six-week post-surgery time point chosen for this study, which could be after any observable injury-induced adaptations. Studies testing earlier or later time points would bring valuable insights on time-dependent oligodendroglial adaptations to chronic pain and inform the best approaches to use for reversing these consequences of neuropathic injury. To this end, in a model of spinal cord injury where OPC are affected as early as 24 hours following injury (Grossman et al., 2001), transplantation of OPC was shown to alleviate pain hypersensitivity (Tao et al., 2013; Yang et al., 2020). At later time points when OPC pools are restored to physiological levels but OL number remain low, strategies targeting OPC differentiation and myelination stimulation may be more appropriate. Remarkably, such an approach alleviated pain-induced cognitive deficits in mice (Zhu et al., 2024). Recently, in naïve mice, noradrenaline was shown to promote OPC differentiation into OL (Fiore et al., 2023; Lu et al., 2023). If this mechanism is maintained in chronic pain conditions, drugs acting on the noradrenergic system, an important pain control system itself (Alba-Delgado et al., 2013; Camarena-Delgado et al., 2022; Gu et al., 2023; Hirschberg et al., 2017; Kimura et al., 2016; Kucharczyk et al., 2022; Llorca-Torralba et al., 2022; Lubejko et al., 2024; Norris, Kuo, et al., 2024), could be repurposed for promoting OPC differentiation. Similarly, repeated Transcranial Magnetic Stimulation (rTMS) can promote myelination (Fabres et al., 2025; Yang et al., 2020) and is under investigation for treating chronic pain (Fernandes et al., 2022; Yang and Chang, 2020). The region-specificity we observed in our study suggests rTMS interventions targeting brain areas with decreased OL counts may have therapeutic potential for chronic pain.

Altogether, we show that chronic neuropathic injury induces long-term alterations in OL density with region- and possibly sex-specific effects in mice. This work further supports the need for a better understanding oligodendroglial cells in the pathophysiology of chronic pain and to test interventions aimed at reversing pain-induced effects on the oligodendrocyte lineage.

## Supporting information

Supplemental Datasheet 1

## Author contribution

L.J.B. and J.G.M. conceived the project. L.J.B., R.O., and G.B. performed the mouse experiments, L.J.B. and R.O. performed the investigation and analyzed the data. L.J.B. and J.G.M wrote the manuscript. L.J.B., R.O., and J.G.M edited the manuscript. J.G.M. acquired funding, provided research supervision and led overall project administration.

## Data Availability

All data presented in this manuscript is available in **Supplemental Datasheet 1**.

## Acknowledgment

We thank the other members of the Al-Hasani and McCall labs for helpful feedback on this project and the Pradhan lab for allowing us to use their microscope. This work was financially supported by the National Institutes of Health (NIH; R01NS135401, J.G.M.; T34GM141639, G.B.) and the ASPIRE program at Washington University in Saint Louis. This manuscript is the result of funding in whole or in part by the NIH. It is subject to the NIH Public Access Policy. Through acceptance of this federal funding, NIH has been given a right to make this manuscript publicly available in PubMed Central upon the Official Date of Publication, as defined by NIH. Biorender.com was used for figure cartoons. We would like to acknowledge the Osage Nation, Missouria, Illinois Confederacy, and many other tribes as the ancestral, traditional, and contemporary custodians of the land where this work was conducted.

---

**Figure S1.**
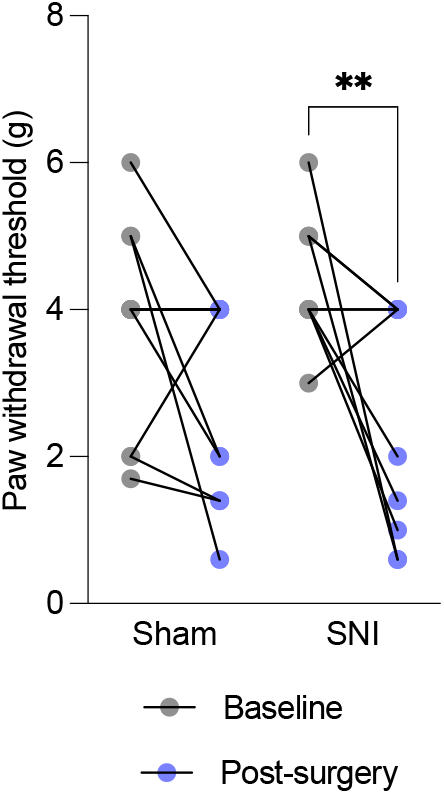
Spared nerve injury (SNI) induces contralateral hypersensitivity. Five weeks after surgery, SNI but not sham mice show a decrease in paw withdrawal thresholds in the hind paw contralateral to the lesion (n_ctrl_ = 11/n_SNI_ = 10; circle points = males, triangle = females.

**Figure S2.**
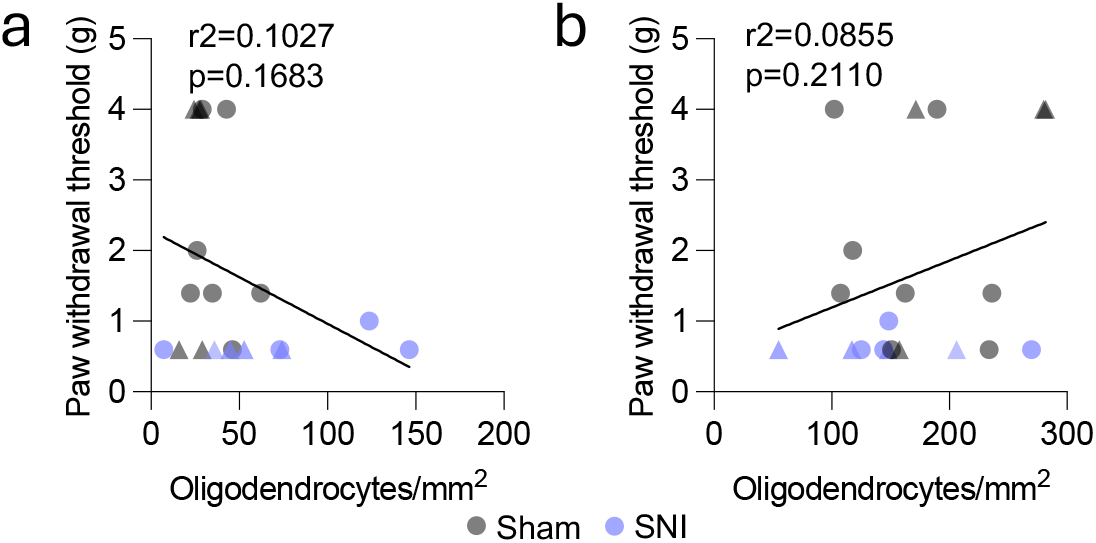
Correlations between OL density and mechanical hypersensitivity in the BLA and PAG. In the BLA (**a**) and the PAG (**b**) the number of oligodendrocytes does not correlate with mice paw withdrawal threshold in the von Frey test.

